# Dynamic binding of the bacterial chaperone Trigger factor to translating ribosomes

**DOI:** 10.1101/2024.03.27.586930

**Authors:** Tora Hävermark, Mikhail Metelev, Erik Lundin, Ivan L. Volkov, Magnus Johansson

**Affiliations:** Department of Cell & Molecular Biology, Uppsala University

## Abstract

The bacterial chaperone Trigger factor (TF) binds to ribosome-nascent chain complexes (RNCs) and co-translationally aids the folding of proteins in bacteria. Decades of studies have given a broad, but often conflicting, description of the substrate specificity of TF, its RNC-binding dynamics, and competition with other RNC-binding factors, such as the Signal Recognition Particle (SRP). Previous RNC-binding kinetics experiments were conducted on stalled RNCs in reconstituted systems, and consequently, may not represent the interaction of TF with ribosomes translating mRNA in the cytoplasm of the cell. Here, we used single-particle tracking (SPT) to measure TF binding to actively translating ribosomes inside living *Escherichia coli*. In cells, TF displays two distinct binding modes — long (ca 1 s) target-specific RNC binding, and shorter (ca 50 ms) sampling of non-target RNCs. RNC binding events are interrupted only by transient excursions to a freely diffusing state (ca 40 ms). We also show that TF competes with SRP for RNC binding *in vivo*, and in doing so, tunes the binding selectivity of SRP.

## Introduction

The nascent polypeptide, emerging at the tunnel exit of the ribosome, is the potential substrate for a large number of targeting factors, modifying enzymes, and chaperones. Depending on the final destination and folding pathway of the mature protein, one or several of these will bind to the ribosome-nascent chain complex (RNC)^1^. Trigger factor (TF) is an ATP-independent chaperone that binds to RNCs and co-translationally aids *de novo* folding of the nascent chain. Once the polypeptide is released from the ribosome, it may stay associated with TF^2,3^, get handed over to other cytosolic chaperones^4,5^, or fold independently^6^. TF has a large substrate pool including cytosolic and secretory proteins, with a preference for sequences of positively charged amino acids flanked by aromatic residues^7^, and is particularly enriched on ribosomes translating outer membrane proteins^8^. In contrast, TF is underrepresented on ribosomes translating inner membrane proteins^8^, which instead are targeted by SRP to the SecYEG translocon for co-translational insertion into the inner membrane^9^. SRP and TF share a docking site on the ribosome close to the polypeptide exit tunnel^10–12^. Whether they compete for binding or can bind simultaneously to the same ribosome has remained elusive as there are studies supporting both cases^13–18^.

TF binds to the ribosome via its N-terminal signature GFRxGxxP motif^12^. The C-terminal domain forms the center of the structure with two protruding arms and facilitates most interactions with the nascent chain^10,19–21^. The middle domain has peptidyl-prolyl isomerase activity as shown by *in vitro* experiments^18,20,21^. The affinity of TF to RNCs is substrate-dependent and recognition of a target nascent chain increases the affinity up to 30-fold compared to a non-target RNC^13,22,23^. However, the times reported for target binding vary greatly, from 1 s to 50 s^13,22–24^. Furthermore, these experiments were done in reconstituted systems with ribosomes stalled on the mRNAs, and consequently may not reflect the kinetics of TF interacting with a growing nascent chain. In a previous single-particle tracking (SPT) study, the TF-RNC binding time *in vivo* was estimated to approximately 0.2 s, but the authors acknowledged that this time represents an average of binding to target RNCs as well as unspecific binding to non-targets^25^.

In this work, we have performed SPT of TF in live *Escherichia coli* at a sufficiently high temporal resolution to distinguish long interactions with target RNCs from short unspecific bindings with non-targets. Moreover, our labeling approach yields long enough trajectories to capture the transient nature of binding and unbinding within the same trajectory. With this experimental methodology, we establish that RNC binding by TF *in vivo* is more dynamic than reported from *in vitro* studies. Moreover, we show that TF competes with SRP for RNC binding, and in doing so, TF tunes the binding selectivity of SRP.

## Results

### SPT captures RNC-bound and free TF

For SPT of TF in live *E. coli*, we first inserted the gene encoding HaloTag downstream of the gene encoding TF on the chromosome, resulting in a C-terminal fusion (TF-Halo). HaloTag covalently binds chloroalkane ligands^26^ such as the chloroalkane-activated organic fluorescent Janelia-Fluor (JF) dyes^27^. We have previously confirmed that the TF-Halo fusion is functional *in vivo* (see Supplementary Note 10B in reference ^28^). We applied our SPT method previously used for HaloTag-labeled ribosomes in live *E. coli*^29^. In short, cells expressing TF-Halo were grown to exponential phase and then incubated with JFX549. Due to the abundance of TF-Halo, we optimized the concentration of JFX549 to ensure that only a subset of TF-Halo molecules was fluorescent, allowing tracking of a single TF-Halo molecule per cell. Cells were sparsely spread onto an agarose pad prepared with growth media on a microscopy slide (Fig. 1a). The sample was incubated at 37°C for at least 1.5 h before imaging, allowing single cells to divide and form small colonies, thus capturing TF-Halo dynamics in growing cells with active translation.

**Fig. 1.**
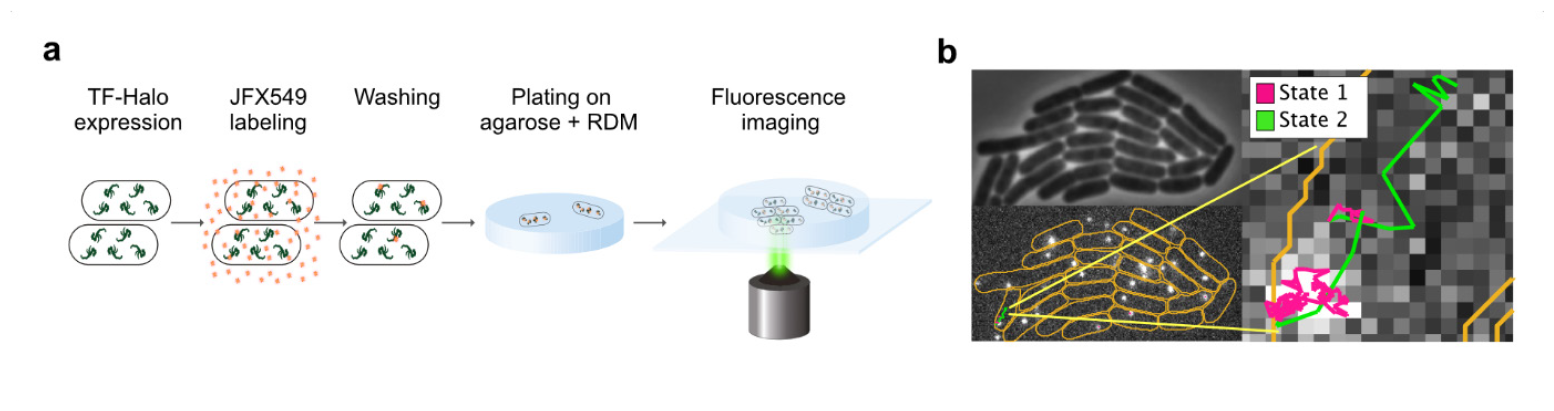
Experimental setup. **a** Cells expressing TF-Halo were incubated with JFX549 and spread on agarose pads containing rich defined medium (RDM) with subsequent incubation at 37°C, allowing formation of small colonies from single cells. Colonies were imaged in phase contrast followed by fluorescence time-lapse movies. **b** Trajectories of single TF-Halo particles were assigned to individual cells (bottom left), segmented based on phase contrast images (top left), and fitted to HMM-based models describing discrete diffusion states (right). The trajectory shown is fitted to a 2-state diffusion model, with steps assigned to state 1 (slow diffusion) and state 2 (fast diffusion) color-coded in magenta and green, respectively. Orange lines are cell outlines.

We performed stroboscopic fluorescence imaging of colonies using a camera exposure time of 5 ms with 3 ms laser illumination per frame. The fluorescence time-lapse movies were analyzed with our pipeline previously used for SPT of tRNA^30^, ribosomal subunits^29^, and SRP^31^. In short, cell outlines were identified based on phase contrast images and a dot detection algorithm identified fluorescing TF-Halo dots in each frame of the movie. Diffusion trajectories of single fluorescent dots were assigned to individual cells, describing the diffusional behavior of the particle over time (Fig. 1b) with a mean trajectory length of 29 frames (Fig. S1).

We observed a distinct diffusional pattern, where the particles were mostly close to immobile and only briefly transitioned into a faster diffusional state (Supplementary Movie 1). Based on our expectations of TF function, we hypothesized that the slow diffusion corresponds to RNC binding and that the faster diffusion is free TF-Halo. To quantify these diffusion differences, we applied a Hidden Markov Modeling (HMM) approach in which all trajectories are fitted to a pre-defined number of discrete diffusion states based on the trajectory step lengths, with state diffusion coefficients and transition frequencies between these states as fitting parameters^29–31^. Each diffusion state, hence, is described by an average diffusion coefficient (D), the steady-state fraction of particles in said state (occupancy), and the average time spent in that state (dwell time).

The HMM algorithm does not provide which model size best fits the data. Since a molecule’s motion in a cell is likely best described by a set of diffusion distributions rather than discrete diffusion states, using statistical tools such as Akaike’s information criterion to find the best model size yield better scores the more states are added (Fig. S2)^29,32^. However, we were interested in finding the simplest model that describes distinct biological functions of TF based on diffusion. We thus fitted the data to models with increasing sizes, ranging from 2-9 diffusion states (Fig. 2a, Supplementary Data 1-4)^29^, and found that three clusters of states appeared independently of model size: one cluster of ribosome-like diffusion states^29,33^ in the range 0.02-0.2 µm^2^s^-1^, a second cluster with tenfold faster diffusion ranging from 2-5 µm^2^s^-1^, potentially corresponding to freely diffusing TF-Halo, and a third cluster containing low-occupancy states (< 1%) with diffusion rates too high to be physiologically relevant (> 10 µm^2^s^-1^). We assign the third cluster to artifacts of the trajectory building, also seen in the analysis of simulated microscopy data^30^.

**Fig. 2.**
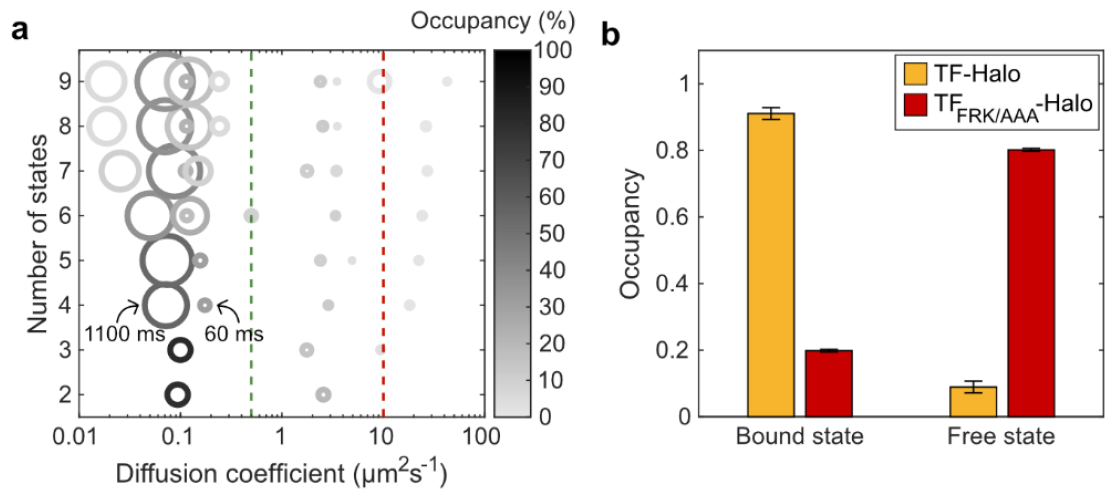
SPT distinguishes free and RNC-bound TF-Halo. **a** HMM fitting of trajectories to different model sizes (2-9). Each model is resolved along the y-axis with discrete diffusion states represented by circles resolved along the x-axis based on diffusion rate. Circles are color-coded according to state occupancy and the area is proportional to the state dwell time. As reference points, two dwell times in the 4-state model are explicitly indicated. Green and red lines mark diffusion thresholds of 0.5 and 10 µm^2^s^-1^, respectively, separating the three clusters of diffusion states (RNC binding, free TF-Halo and tracking artefacts). Data shows chromosomally expressed TF-Halo, n = 99,352 trajectory steps cumulated from 3 independent experiments. Full HMM output is found in Supplementary Data 4 **b** Occupancy in 2-state models of TF-Halo and TF_FRK/AAA_-Halo, expressed from plasmids in a TF knockout strain. The 2-state models are weighted averages of models with 4-9 states coarse-grained into two states using a threshold of 0.5 µm^2^s^-1^ to separate RNC binding from free TF-Halo. n = 83,546 and 75,585 trajectory steps from 3 and 5 independent experiments for TF-Halo and TF_FRK/AAA_-Halo, respectively. Error bars represent the weighted standard deviation calculated from coarse-grained models of 4-9 states. Full output of coarse-grained models is shown in Supplementary Data 5 and output from all model sizes is shown in Supplementary Data 6-7.

To specifically investigate RNC binding events, we coarse-grained the larger HMM-fitted models into 2 states using a threshold at 0.5 µm^2^s^-1 29–31,33^, with state 1 corresponding to RNC-bound TF-Halo and state 2 to free TF-Halo. The occupancy and average dwell time, calculated as weighted averages from course-grained models 4-9, were 85% and 180 ms for the bound state, and 15% and 28 ms for the free state, respectively. To verify that the slow diffusion state captures RNC binding, we tracked a TF mutant in which residues 44-47 (FRK) were exchanged to AAA, thus compromising ribosome binding^12,25^. Indeed, TF_FRK/AAA_-Halo displayed a slow-state occupancy of only 20%, confirming that slow diffusion corresponds to RNC binding by TF (Fig. 2b, Fig. S3, Supplementary Movie 2).

### RNC binding kinetics depend on the cellular level of TF-Halo

The RNC-bound dwell time suggested in the 2-state model, 180 ms, likely represents an average of both unspecific and specific binding to RNCs, in line with previous *in vivo* TF SPT results^25^. To investigate if the average RNC-bound time could be perturbed, we titrated the TF-Halo levels in a TF knockout strain (Fig. 3, Supplementary Data 5 and 7-11). With gradual increase of TF-Halo, the RNC-bound dwell time decreases significantly, with proportionally less effect on the RNC-bound occupancy, supporting that the average dwell time in this 2-state model is a combination of longer, target RNC bindings, and shorter samplings. At higher expression levels, the system becomes saturated also with respect to unspecific binding, as reflected by a drastic decrease also in the RNC-bound occupancy.

**Fig. 3.**
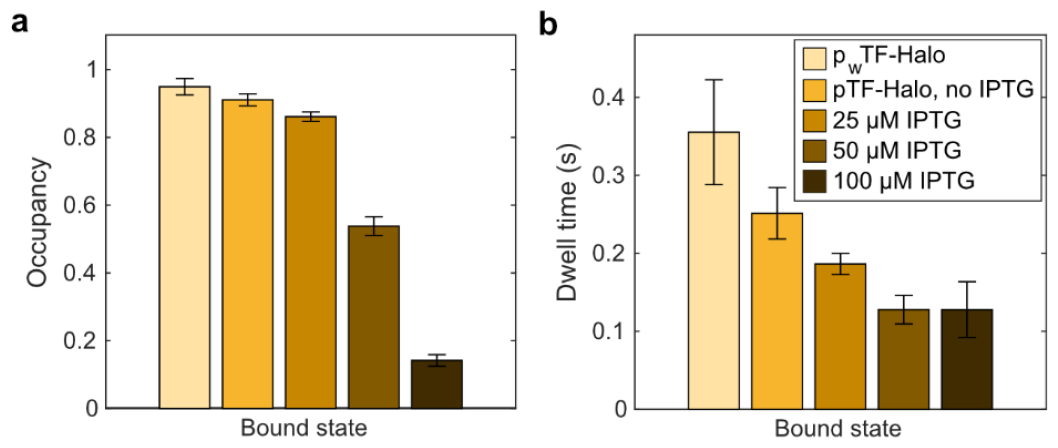
Increasing TF-Halo levels reduce the occupancy and average dwell time of RNC binding. Occupancy (**a**) and dwell time (**b**) in the RNC-bound state in 2-state models. TF-Halo was expressed from plasmids in a TF knockout strain, either with weak constitutive (p_w_TF-Halo) or IPTG-inducible expression (pTF-Halo with 0, 25, 50 and 100 µM IPTG). The 2-state models are weighted averages of models with 4-9 states coarse-grained into two states using a threshold of 0.5 µm^2^s^-1^. N = 110,271; 83,546; 107,952; 86,308; 62,507 steps from 3, 3, 3, 3, and 4 independent experiments for each condition as listed in the figure legend. Error bars represent the weighted standard deviation calculated from coarse-grained models of 4-9 states. Full output from coarse-grained models is shown in Supplementary Data 5 and output from all model sizes is shown in Supplementary Data 7-11.

These titration experiments highlight that the kinetic parameters in our 2-state models depend on the number of TF molecules competing for RNC binding. In order to investigate whether the stoichiometry between RNCs and TF-Halo in the chromosomally tagged strain is representative of the native situation, we also performed SPT of TF-Halo expressed in low copy number (Fig. S4) from a plasmid in a wild-type (wt) TF background, as well as in the chromosomal TF-Halo background. We find that the RNC-bound state occupancy of TF-Halo is slightly lower when competing with wt TF (73%) than when competing with chromosomally expressed additional TF-Halo (83%, Fig. 4a, Supplementary Data 12-13). This result suggests that the cellular level of TF-Halo is lower than that of wt TF when they are expressed from the same position on the chromosome, or, alternatively, that TF-Halo has slightly reduced binding capacity to RNCs compared to the wt TF. However, since TF-Halo is functional^28^, and since the overall RNC binding dynamics of TF-Halo is highly similar in all three scenarios (Fig. 4b, average bound-state dwell times 130-180 ms, and free-state dwell time 28-43 ms), we believe that our measurements represent the binding kinetics of TF to RNCs well. The parameters measured from low-level TF-Halo in the wt TF background (Fig. 4c), though, are likely the most accurate.

**Fig. 4.**
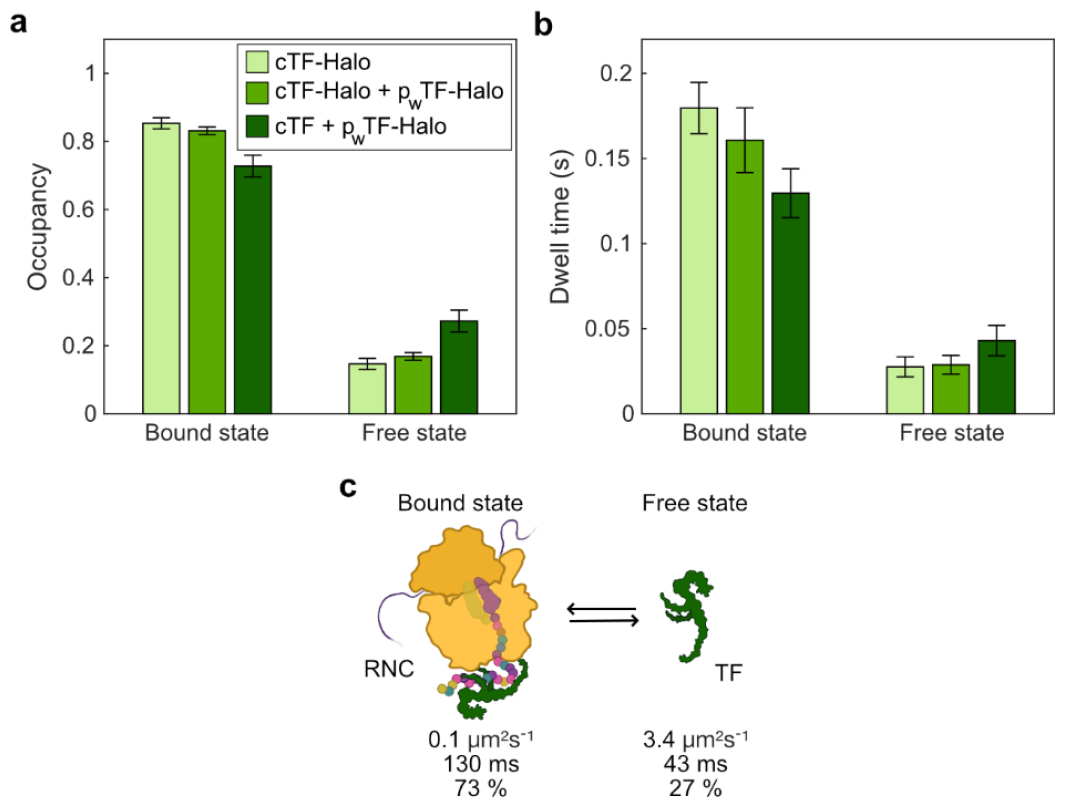
2-state models of TF-Halo in different background strains. Occupancy (**a**) and dwell time (**b**) in 2-state models from SPT of TF-Halo in the chromosomally tagged strain (cTF-Halo), the chromosomally tagged strain with additional weak plasmid expression (cTF-Halo + p_w_TF-Halo) and in a wt strain with additional weak plasmid expression (cTF + p_w_TF-Halo). **c** 2-state model based on cTF + p_w_TF-Halo. The 2-state models are weighted averages of models with 4-9 states coarse-grained into 2 states using a threshold of 0.5 µm^2^s^-1^. N = 99,352; 92,876; and 96,321 trajectory steps cumulated from 3, 3 and 4 independent experiments for cTF-Halo, cTF-Halo + p_w_TF-Halo and cTF + p_w_TF-Halo, respectively. Error bars represent the weighted standard deviation calculated from coarse-grained models of 4-9 states. Full output from coarse-grained models is shown in Supplementary Data 5 and output from all model sizes is shown in Supplementary Data 4, 12 and 13.

### Larger models resolve RNC sampling from target binding

From the coarse-grained 2-state analysis of TF-Halo diffusion, we established that TF is RNC-bound 73% of the time with an average dwell time of 130 ms (Fig. 4c, low TF-Halo expression in a wt background), which we assume is a global average of target binding and shorter, unspecific bindings. However, in all fitted models with 4 states or more, we robustly identified one RNC-bound state with significant occupancy (around 30%) and approximately 20-fold shorter average dwell time than the other RNC-bound states (as shown by the differences in circle areas in the bubble plot, see Fig. 2a for chromosomally expressed TF-Halo and Fig. S5 for low TF-Halo in a wt background). We speculate that these long- and short-lived RNC-bound states are separated in the HMM analysis not due to differences in diffusion rates, but rather, different underlying binding kinetics between sampling and target binding. Since the 4-state model is the simplest one where the potential sampling state appears, both for chromosomally expressed TF-Halo (Fig. 2a, Supplementary Data 4) and for low TF-Halo expression in a wt background (Fig. S5, Supplementary Data 13), we investigated this model further. For low TF-Halo expression in a wt background, state 4 is fast diffusing (D_4_ = 12 µm^2^s^-1^) with 2% occupancy, corresponding to the aforementioned tracking artefacts, and is, hence, disregarded from the biological interpretation. We tentatively assign free TF-Halo to state 3 (D_3_ = 3 µm^2^s^-1^), RNC sampling to state 2 (D_2_ = 0.2 µm^2^s^-1^), and target RNC binding to state 1 (D_1_ = 0.1 µm^2^s^-1^). According to this model, 48% of all TF-Halo molecules display target binding at steady state with an average dwell time of approximately 800 ms (Fig. 5a, Supplementary Data 13). The occupancy and dwell time in the sampling state is 27% and 54 ms, respectively, whereas 23% of TF-Halo is in the free state where it stays for, on average, 42 ms before binding a new ribosome. By inspecting the HMM estimated fluxes of molecules between states, we find frequent transitions between states 2 and 3 (as exemplified in Fig. 5b, Fig. S6) with occasional long-lasting excursions to state 1 (Fig. 5c, Fig. S6), also apparent by eye in the fluorescence movies (Supplementary Movie 3 and 4). This pattern indicates that TF-Halo is exerting a sampling-like behavior, scanning for target RNCs in the time range of tens of milliseconds, and that this is occasionally (one per 20 sampling events) interrupted by more long-lasting (second range) interactions with target RNCs. The same pattern is seen with the chromosomally expressed TF-Halo, with two RNC-bound states of 1100 ms and 59 ms, respectively (Fig. 2a, Fig. S6, Supplementary Data 4). In contrast, trajectories of TF_FRK/AAA_-Halo, compromised in ribosome binding, show fast diffusion without any such frequent transitions between states 2 and 3 (Fig. 5d, Supplementary Movie 5), confirming that state 2 captures short samplings of ribosomes rather than any other slow-diffusing, ribosome-independent, activity of TF-Halo.

**Fig. 5.**
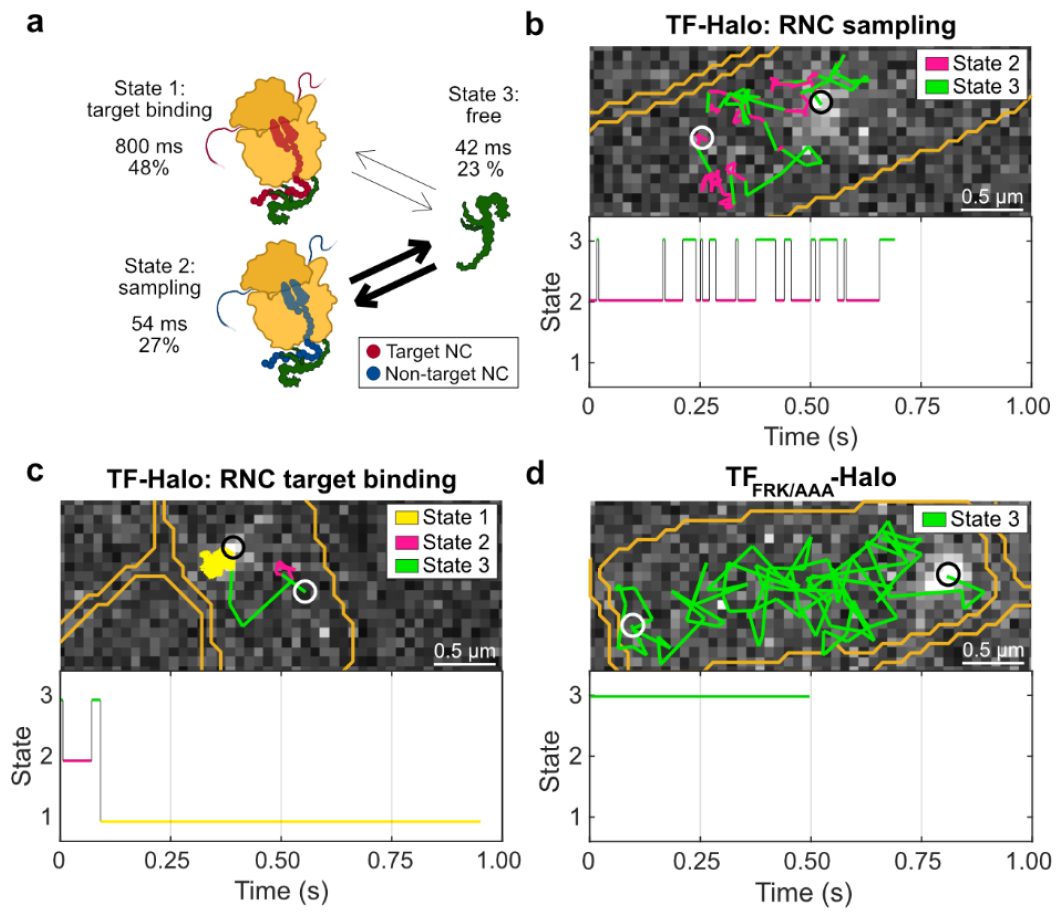
Larger models resolve stable TF-RNC binding from sampling. **a** Biological interpretation of 4-state model. The thickness of the arrows is proportional to the relative fluxes of particles between states. State 4, with D = 12 µm^2^s^-1^ and 2% occupancy, is assigned to tracking artefacts and thus not included in the biological interpretation. The complete model with state 4 included is shown in Fig. S7 and Supplementary Data 13. The model is based on SPT of low levels of TF-Halo in a wt TF background (cTF + p_w_TF-Halo), n = 96,321 trajectory steps cumulated from 4 independent experiments. A similar 4-state model was obtained from chromosomally expressed TF-Halo (Supplementary Data 4). **b** Example of TF-Halo trajectory displaying short RNC samplings. **c** Example of a TF-Halo trajectory displaying long RNC target binding. **d** Example trajectory of TF_FRK/AAA_-Halo. Trajectories in **b-d** are HMM-fitted to 4-state models The top panels display trajectories in space and the bottom panels show the state transitions in each trajectory over time. Steps are color-coded according to state assignments, with states 1, 2 and 3 corresponding to RNC target binding, sampling, and free TF-Halo, respectively. White and black circles indicate start and end points of a trajectory, respectively. Cell outlines are shown in orange.

Further, we confirmed that this model is robust with respect to time resolution by reducing the camera exposure time from 5 ms to 2.5 ms. With 2.5 ms exposures, the distinction between sampling and target RNC-binding remained (Fig. S8, Supplementary Data 14). However, by increasing the exposure time to 60 ms (as in the TF SPT in reference ^25^), the distinction disappeared (Fig. S8, Supplementary Data 15), highlighting why a temporal resolution higher than the estimated sampling time is necessary for distinguishing the different binding kinetics.

When comparing the kinetic models of TF-Halo and the FRK/AAA mutant, we noticed that by removing the ribosome binding ability, and consequently, the fast transitions between free TF-Halo and RNC sampling, the free state dwell time in the 2-state models increased by 100-fold, from tens of milliseconds to seconds (Fig. 6). Thus, the free state dwell time is an indicator of whether TF is sampling RNCs or not. With this observation in mind, we revisited the TF-Halo titration experiments (Fig. 3), where we concluded that the more TF-Halo molecules there are in a cell, the more RNC sampling in relation to functional binding there will be. Inducing TF-Halo expression with 25 µM Isopropyl β-D-1-thiogalactopyranoside (IPTG) did not affect the free state dwell time, indicating that there is substantial sampling occurring (Fig. 6). When inducing TF-Halo expression with 50 and 100 µM IPTG, the dwell time increased to 89 ± 20 and 660 ± 170 ms, respectively, confirming our previous conclusion that at such high expression levels, there is competition not only for stable binding, but also for RNC sampling.

**Fig. 6.**
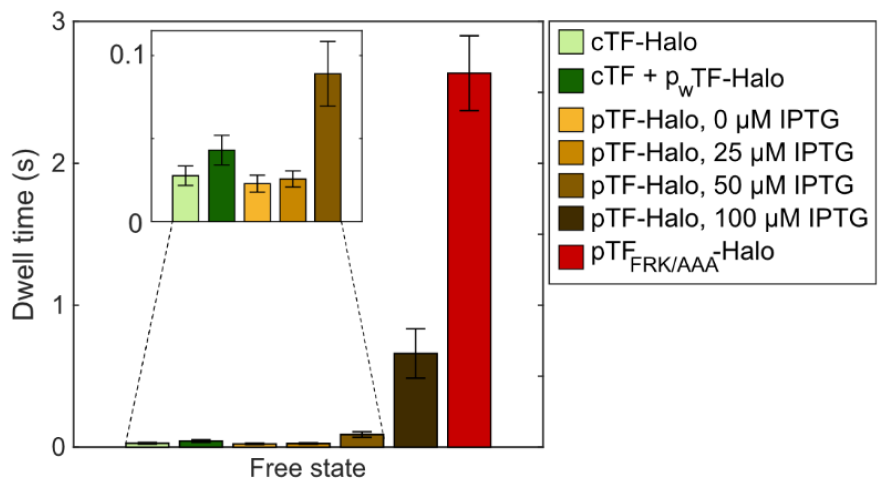
Free state dwell times of TF-Halo under different experimental conditions. Free state dwell times in 2-state models obtained from weighted averages of coarse-grained models (4-9) with a threshold of 0.5 µm^2^s^-1^. Conditions with the prefix “c” encode chromosomal expression of TF or TF-Halo, “p_w_” encodes low constitutive expression of TF-Halo from a plasmid and “p” is IPTG-inducible plasmid expression in a TF knockout strain. n = 99,352; 96,321; 83,546; 107,592; 86,308; 62,507; and 75,585 trajectory steps from 3, 4, 3, 3, 3, 4, and 5 independent experiments for each condition as listed in the figure legend. All coarse-grained models are shown in Supplementary Data 5 and full HMM models in Supplementary Data 4, 6-10 and 12. Error bars represent weighted standard deviation calculated from the included model sizes (4-9).

### A nascent polypeptide chain is needed for TF binding to ribosomes

Previous *in vitro* studies have shown that TF displays weak-affinity binding to vacant, non-translating, 70S ribosomes, with a dissociation rate similar as for RNCs displaying non-target nascent chains^13,23^. To our knowledge, the binding kinetics of TF to free 50S subunits have not been measured *in vitro*, but stable TF binding to free 50S subunits have been captured for structural analyses^10,34,35^. Thus, in the previous *in vivo* TF SPT study^25^, they concluded that interactions of TF with free 50S subunits were included in the diffusion model. To investigate the nature of the observed TF-RNC sampling state, we treated cells with Kasugamycin (Ksg), an inhibitor of translation initiation^36^. Our previous SPT of ribosomal subunits showed that, under identical Ksg treatment conditions, 80% of the 50S subunits are freely diffusing^29^. Hence, by tracking TF-Halo in Ksg treated cells, we expected a higher proportion of short samplings relative to target RNC bindings. However, upon Ksg treatment, the overall bound-state occupancy decreased from 85 ± 2% to 40 ± 6% (Fig. 7a), whereas the bound-state dwell time increased almost twofold (from 180 ± 20 ms to 330 ± 40 ms, Fig. 7b). These results indicate a smaller proportion of samplings to functional bindings, rather than the expected opposite, implying that TF-Halo binds to free 50S subunits less efficiently than to elongating 70S. In support of this hypothesis, the free-state dwell time also increased drastically from 28 ± 6 ms to 500 ± 160 ms (Fig. 7b), similar to what was observed for TF-Halo overexpression (Fig. 6, induction with 100 µM IPTG), indicating that TF-Halo is freely diffusing for a longer time before finding a vacant binding site. If TF was to sample free 50S subunits equally well as elongating 70S ribosomes, we would not observe such an effect, as the number of possible binding sites, whether they be in 50S or 70S configuration, would be constant. Finally, by inspecting the individual HMM estimated models of different sizes, we also note that the sampling state, apparent in all model sizes with 4 states or more in the absence of Ksg (Fig. 2a, Fig. S5), now only shows up in 7-state models or higher, with an occupancy of only around 6%, compared to roughly 30% in untreated cells (Fig S9, Supplementary Data 4 and 16). Hence, we conclude that the transient (tens of milliseconds) RNC-bound state observed in our experiments, represent samplings of translating ribosomes, likely synthesizing non-TF-target polypeptides, and that sampling events of non-translating 50S subunits, if such events occur, must be faster than the temporal resolution of our experiments, i.e. less than a few milliseconds.

**Fig. 7.**
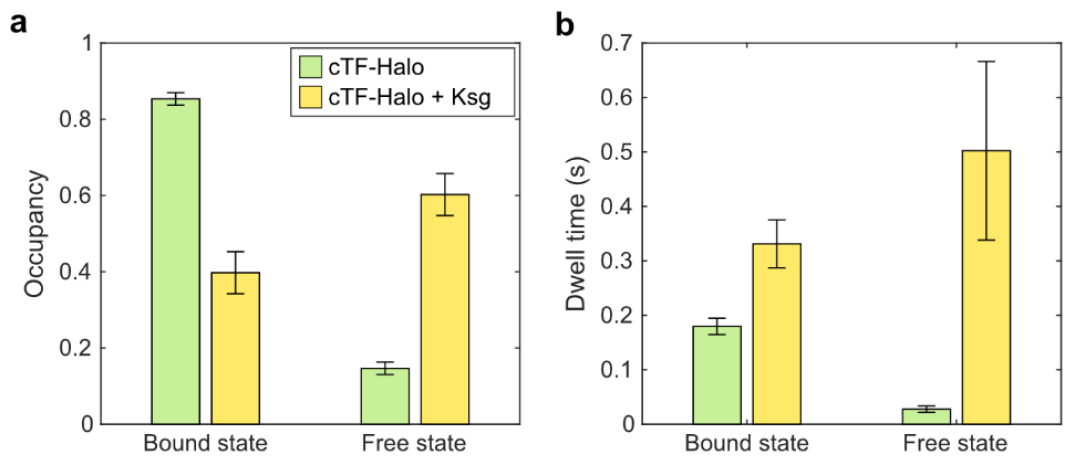
TF-Halo in cells treated with Kasugamycin. Occupancy (**a**) and dwell time (**b**) in 2-state models from SPT of TF-Halo in the chromosomally tagged strain (cTF-Halo), with and without 2 mg/ml Ksg added to the sample. The 2-state models are weighted averages of models with 4-9 states coarse-grained into 2 states using a threshold of 0.8 µm^2^s^-1^. The threshold was determined based on the diffusion rate of 50S subunits in Ksg-treated cells^29^ to separate TF-Halo bound to 50S and 70S ribosomes from free TF-Halo. N = 99,352 and 85,146 trajectory steps cumulated from 3 independent experiments each. Error bars represent the weighted standard deviation calculated from coarse-grained models of 4-9 states. Full HMM outputs from all model sizes are shown in Supplementary Data 4 and 16, and coarse-grained models are shown in Supplementary Data 5.

### TF-Halo and SRP compete for RNC-binding *in vivo*

We next sought to investigate the possible competition for RNC binding between TF and SRP. Since the cellular levels of TF are much higher than those of SRP (40-90 µM TF and approximately 1 µM SRP^37–41^) we assumed that increasing the SRP levels would not yield a detectable effect on TF-Halo activity. Instead, we applied our protocol for SRP tracking^31^ and explored the effect on SRP-RNC binding upon titration of TF-Halo, induced by IPTG, in a TF knockout strain. With increasing TF-Halo concentration, the RNC-bound fraction of SRP decreased, from 46.3 ± 0.4% at the lowest TF-Halo concentration, to 26.8 ± 0.2% at the highest (Fig. 8a Supplementary Data 17-20). The effect is TF-specific, as no such reduction was seen when HaloTag alone was titrated using the same expression system (Fig. 8c, Supplementary Data 21 and 22). Thus, we conclude that TF and SRP compete for ribosome binding *in vivo*.

**Fig. 8.**
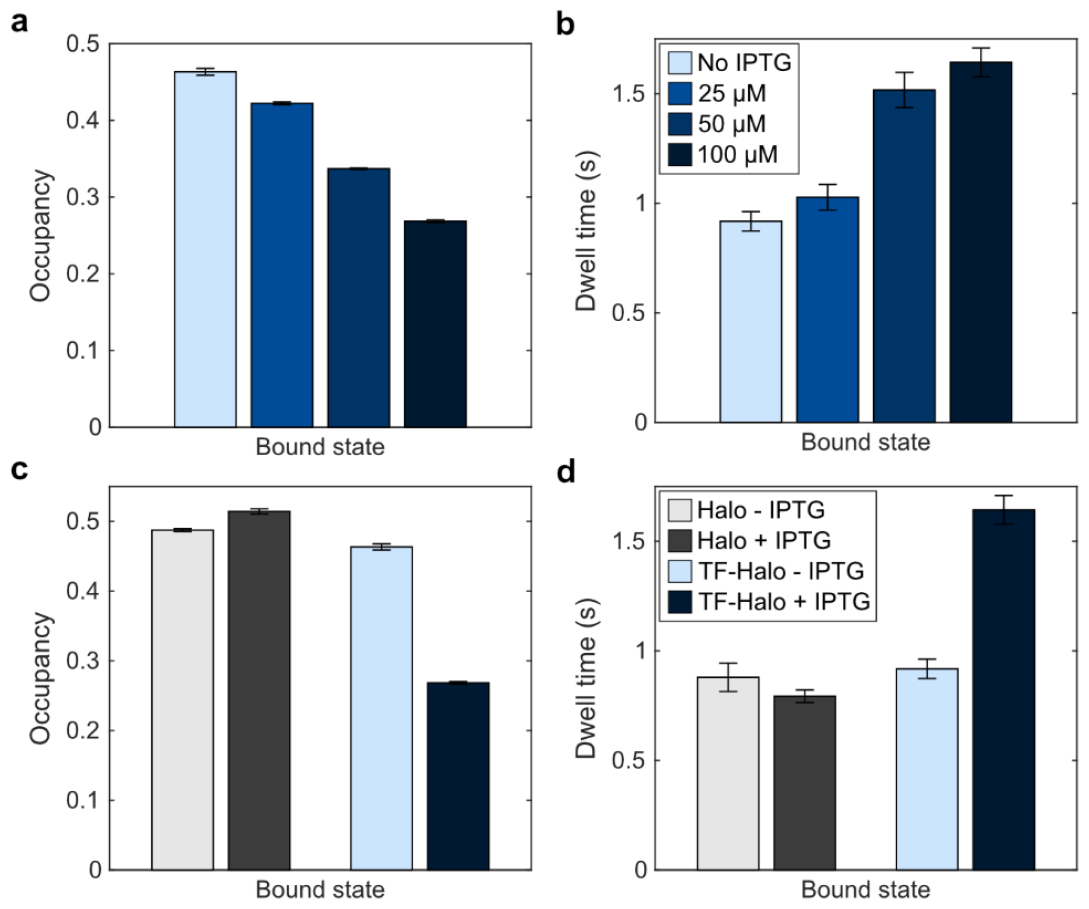
TF-Halo competes with SRP for RNC binding. Occupancy (**a**) and dwell time (**b**) of SRP in the RNC-bound state when inducing expression of (dark) TF-Halo with 0, 25, 50 and 100 µM IPTG from a plasmid in a TF knockout strain. **c**,**d** Occupancy and dwell time of SRP in the RNC-bound state with 0 and 100 µM IPTG induction of either HaloTag alone or TF-Halo. Data show 2-state models obtained by weighted averages of models with 4-9 states coarse-grained into two states using a threshold of 0.5 µm^2^s^-1^. n = 79,881; 65,506; 54,544; 66,430; 81,499; 106,826 trajectory steps for induction of TF-Halo with 0, 25, 50 and 100 µM and 0 and 100 µM of HaloTag, respectively. Each condition was tested in 3 independent experiments. Error bars represent weighted standard deviation calculated from the included model sizes (4-9). Output from coarse-grained models is shown in Supplementary Data 5 and output from all model sizes is shown in Supplementary Data 17-22.

Since TF and SRP have distinct substrate pools, i.e., SRP mainly targets ribosomes translating inner membrane proteins whereas TF targets cytosolic and secretory proteins^8,9^, we first hypothesized that the reduction in RNC binding by SRP is a consequence of increased unspecific binding by TF-Halo, making those RNCs which are translating SRP targets less accessible to SRP. However, we observe that the dwell time of SRP on RNCs gradually increases with elevated TF-Halo levels (Fig. 8b), suggesting that at low TF levels, there is a larger fraction of unproductive SRP-RNC interactions, which consequently reduces the average dwell time. Thus, a more likely explanation to the observed reduction in RNC binding by SRP would be that, at low TF-Halo concentration, SRP also binds TF targets to a substantial extent. Again, the effect is TF specific, since the dwell-time of SRP on ribosomes does not change significantly upon overexpression of HaloTag alone (Fig. 8d). Hence, our results imply that in the absence of TF-Halo, SRP displays more unproductive RNC bindings, suggesting that the large number of TF molecules in a normal cell actually tune the selectivity of SRP.

## Discussion

TF is an extensively studied chaperone, but there is a lack of consensus regarding its RNC-binding properties, and we have had limited knowledge of its overall activity in the cell. In this work, we report TF binding to translating ribosomes in living cells at a high enough temporal resolution to distinguish unspecific binding of non-target RNCs from more stable binding to targets. Although kinetics experiments in reconstituted systems have suggested that such differences between target binding and sampling exist, this work is the first to capture them in the native environment, and during ongoing protein synthesis. We find that TF mainly samples RNCs in search of target nascent chains. The average sampling time is around 50 ms and stable binding is on the one-second time scale (800 ms for TF-Halo expressed at low levels in a wt TF background and 1000 ms for TF-Halo expressed from the chromosome). These dwell times contradict early *in vitro* data^22,23^, but are in good agreement with a more recent study by Bornemann et al., reporting average dwell times on non-target RNCs of 80 ms (Lep75-RNCs) and target RNCs of 2.5 and 1.6 s (proOmpA75-RNC and HemK75-RNC)^13^. Whereas these *in vitro* studies suggested similar binding to non-translating 70S ribosomes as to non-target RNC’s, however, our TF SPT in Ksg treated cells suggest that a protruding peptide is needed for ribosome binding. It should also be noted, though, that those *in vitro* numbers represent binding to stalled RNCs using a reconstituted system, so a direct comparison is perhaps not completely relevant.

We acknowledge that our experimental approach captures the average target binding kinetics to all kinds of target RNC substrates, which may vary depending on the amino acid sequence and length of the nascent chain^13,22^. We also recognize that exact dwell times are to some degree model-dependent. Nevertheless, we robustly see that functional binding lasts for on average between 0.8-2 s, and sampling 40-70 ms, independent of model size and expression system (cTF-Halo or p_w_TF-Halo in wt background, Supplementary Data 4 and 13, respectively), supporting a transient binding and re-binding cycle. In contrast to in vitro-based RNC-binding kinetics data, selective Ribo-Seq of TF indicates that binding and unbinding is a dynamic process, possibly with many binding events per translation event, as the coverage of TF on transcripts is not uniform^8^. Since our data support a model in which TF and SRP competes for RNC binding, we pose that these transient binding dynamics better explain the role of TF as a promiscuous chaperone with a large substrate pool overlapping with other processing factors^13,42^. If TF would stay bound to the majority of RNCs for up to 50 s^23^, this would in practice yield most RNCs inaccessible to other essential processing factors, which would be detrimental to the cell. Promiscuous, but dynamic, binding to practically all potential targets is likely a general feature of biological systems, but has been difficult to study with conventional binding assays since it requires in situ studies at very high time resolution.

TF dimerizes with an apparent K_D_ between 1-18 µM and a half-life of 1 s *in vitro*, but it binds to ribosomes as a monomer^2,43^. Assuming that the cellular concentration is at least 40 µM^37,39,40^ and with a free state occupancy of 27% (Fig. 5a, Supplementary Data 13), we thus expect the concentration of free TF to be around 11 µM in our experiment, i.e., high enough for dimerization to occur at a significant level *in vivo*^43,44^. Strikingly, however, our SPT data show that the average time between RNC-binding events is only tens of milliseconds. Hence, with a half-life of 1 s for the dimer, we deem it unlikely that TF dimerizes between RNC-binding cycles *in vivo*. This finding further highlights the significance of single-particle experiments in exploring the dynamic nature of biomolecules.

With this work, we have provided an experimental scheme to study co-translational processing of nascent chains in its native environment. We have directly measured both sampling and functional binding of TF to active ribosomes, and by genetic perturbations, measured the RNC-binding competition between SRP and TF. This method can further be used for measuring the binding kinetics of other co-translational processing factors to RNCs, and their dynamic interplay, in living *E. coli* cells.

## Methods

### Cloning and strain construction

Cloning procedures involving PCR were performed using Q5 High-fidelity polymerase (New England Biolabs) according to manufacturer’s protocol. For construction of the chromosomal TF-Halo fusion in *E. coli* K12 MG1655 (accession #U00096), the gene encoding HaloTag was inserted on the chromosome directly upstream of the stop codon of the *tig* gene, encoding TF, creating a C-terminal TF-Halo fusion (MG1655tig::halotagKan). The *halotag* gene was inserted along with a kanamycin resistance marker using lambda red recombineering^45^ with pKD4_halotag^29^ as template and primers tig_HalonIns_F and tig_del_R (Table S1). The tig_HaloIns_F primer was designed such that the start codon of *halotag* is exchanged to GGC, creating a 1-glycine linker between TF and HaloTag.

For experiments with titration of TF-Halo levels, the fusion was expressed from a plasmid (pQE30lacIq_tig_halotag) under the T5 promoter with two lac operators, yielding IPTG-inducible expression. The plasmid was inserted in a strain lacking chromosomal TF (MG1655Δtig::kan). Similarly, the TF_FRK/AAA_-HaloTag mutant was expressed from the same vector (pQE30lacIq_tig_halotag_frk_aaa) in the TF knockout background. Construction of MG1655Δtig::kan and the two plasmids are described in Amselem et al. 2023^28^. For low constitutive expression of TF-Halo from a plasmid (p_w_TF-Halo), inverse PCR was performed on pQE30lacIq_tig_halotag to exchange the T5 promoter to the weak constitutive apFAB124 promoter and to switch to a weak Shine-Dalgarno sequence (TTCTCA instead of AGGAGG^29^) using primers pQE_SDw_F and p124_R (pQE30lacIq_p124_SDw_tig_halotag). The linear PCR product was phosphorylated with T4 Polynucleotide kinase and circularized using T4 DNA-ligase according to manufacturer’s instructions (Thermo Scientific). All constructs were verified by Sanger Sequencing.

### Sample preparation

For SPT of chromosomally expressed TF-Halo, an overnight culture of MG1655tig::halotagKan was prepared from a glycerol stock in Luria Broth (LB) at 37°C 200 rpm. The overnight culture was diluted 1:100 in fresh LB followed by incubation at 37°C 200 rpm until OD_600_ reached 0.4-0.8. Cells were harvested and resuspended in EZ Rich Defined Medium (RDM) with 0.2% glucose (Teknova) and 0.2 µM JFX549 (a gift from Luke Lavis lab). Cells were labelled at 25°C for 30 min, followed by washing with M9 medium with 0.2% glucose, incubation in RDM + glucose at 37°C 200 rpm for 60 min, and three subsequent washing steps in M9 + glucose to remove any unbound JFX549. The culture was diluted to approximately OD_600_ 0.003 and spread onto a pad with 2% agarose (SeaPlaque GTG Agarose, Lonza) in RDM + glucose on a microscopy slide. The sample was put in the microscope surrounded by an incubator set to 37 ± 2°C, and left for incubation for at least 100 min and a maximum of 200 min before imaging. For SPT of TF-Halo in Ksg-treated cells, the sample was incubating for 100 min after which 2 mg/ml Ksg was injected to the sample. Data was collected after 50-80 min Ksg treatment.

SPT of TF-Halo expressed from the pQE30lacIq_tig_halotag plasmid was performed in the MG1655Δtig::kan strain. Sample preparation was performed as described above, with all growth media supplemented with 100 µg/ml ampicillin for maintenance of the plasmid. For induction of TF-Halo expression, IPTG was added to the RDM-agarose mixture during preparation of the agarose pad to a final concentration of either 25, 50 or 100 µM. Sample preparation for tracking the TF_FRK/AAA_-HaloTag mutant in MG1655Δtig::kan was performed similarly, without IPTG induction, yielding only leaky expression of TF_FRK/AAA_-HaloTag.

For tracking of SRP, LD655-labeled 4.5S RNA was electroporated into MG1655Δtig::kan carrying either pQE30lacIq_tig_halotag for IPTG-inducible expression of TF-Halo, or pQE30lacIq_halotag (lab collection) for inducible expression of only HaloTag. Preparation of LD655-4.5S RNA, electrocompetent cells and electroporation was executed as described in Volkov *et al*. (2022)^31^ and IPTG induction on the agarose pad was achieved as described above.

### Optical setup and imaging conditions

For widefield epifluorescence imaging, an inverted Ti2-E microscope (Nikon) with a CFI Plan Apo lambda 1.45/100x objective (Nikon) was used. The microscope was built into an H201-ENCLOSURE incubator hood H201-T-Unit-BL and controller (OKOlab), allowing imaging at 37±2°C. Brightfield and fluorescence images were acquired with an iXon 897 Ultra EMCCD camera (Andor) with an additional 2x lens in front of it (DD20NLT, Diagnostic Instruments). Phase contrast images were acquired with the same iXon camera or with an Infinity 2-5M camera (Lumenera). For imaging of HaloTag-fused proteins labeled with JFX549, either a 553 nm laser (SLIM-553L, 150 mW, Oxxius) or a 546 nm laser (2RU-VFL-P-2000-546-B1R, 2000 mW, MPB Communications) was used with a power density of 3 kW/cm^2^ on the sample plane and stroboscopic illumination with 3 ms laser pulses per 5, 30 or 60 ms camera exposure. JFX549-labeled TF-Halo was imaged at 5 and 60 ms. For TF-Halo imaging using camera exposures of 2.5 ms, 2.3 ms laser pulses was used. For tracking of LD655-labeled 4.5S RNA, a 639 nm laser (Genesis MX 639-1000 STM, Coherent) was used with a power density of 4.5 kW/cm^2^ on the sample plane and stroboscopic illumination with 1.5 ms laser exposure per 20 ms camera exposure. Between 500-1000 fluorescence images were acquired per cell colony to create the fluorescence movies. To visualize SYTOX blue stain on cells electroporated with LD655-4.5S RNA, and thus identify and exclude any dead cells, they were imaged with a 405 nm laser (06-MLD 360mW, Cobolt) with a power density of 17 W/cm^2^ and continuous illumination with 21 ms camera exposure time.

The microscope was controlled using µManager and data acquisition was automated using custom-made µManager plugins. Between 30-90 positions on the agarose pad were recorded in one experiment, with each position containing mini colonies grown from single cells. Each tracking condition was repeated in at least three independent experiments, showing consistent results between replicates, see model outputs from each replicate and cumulated datasets for chromosomal TF-Halo expression in Supplementary Data 1-4.

## Data analysis

SPT data was analyzed using a custom MATLAB-based pipeline, as previously described^29,31,46^. In short, segmentation masks of single cells were created based on phase contrast images using the Per Object Ellipse fit (POE) method for adaptive thresholding^47^. Segmentation parameters were optimized based on which camera was used for phase-contrast imaging (iXon 897 or Infinity 2-5M) to obtain equivalent segmentation performance. Each position was manually curated to remove any poorly segmented cells and cells that were placed outside the region of even fluorescence illumination within the field of view. For SRP tracking, any SYTOX-positive cells, i.e., cells that died during electroporation, were also removed. Segmentation masks were aligned with fluorescence images for tracking fluorophores on a per-cell basis, using the radial symmetry-based algorithm^48^. Symmetric Gaussian PSF modelling and maximum a posteriori fitting was performed for dot localization refinement and for estimation of dot localization uncertainty to filter out any faulty dots^49^. Dot detection and refinement parameters were optimized for each type of fluorophore (JFX549 and LD655) and laser used (546, 553 and 639 nm). Each detected dot was assigned to a segmented cell and once there was only one dot detected in a cell, the trajectory of said dot was built using the uTrack algorithm^50^.

To extract the diffusional behavior of particles, an ensemble of trajectories with a minimal length of 5 steps (TF-Halo) or 3 steps (SRP) were analyzed using the HMM-based algorithm as previously described^29,31,49^. The trajectories were fitted to a pre-defined number of hidden states, corresponding to discrete states of diffusion with an average diffusion rate, steady state occupancy and dwell time. The data from each tracking experiment was analyzed separately as well as combined with biological replicates, yielding cumulated datasets containing at least 50 000 steps per experimental condition. The data was fitted to model sizes ranging from 2-9 diffusion states. 2-state models were obtained by coarse-graining all larger models (4-9 states) and calculating weighted averages of the fit parameters. For TF, a threshold of 0.5 µm^2^s^-1^ was used during coarse-graining to separate free and ribosome-bound TF. For SRP, 0.8 µm^2^s^-1^ was used to separate ribosome-bound from free SRP (4.5S RNA + Ffh) and free 4.5S RNA (in agreement with reference ^31^).

## Acknowledgements

We are thankful to the Luke Lavis laboratory for providing the JFX549-HaloTag dye, and to Irmeli Barkefors for providing feedback on the text. This work was supported by the European Research Council (M.J. 947747-SMACK), and the Swedish Research Council (M.J. 2019-03714, 2023-03383). Computations and data handling was enabled by resources from the Swedish National Infrastructure for Computing at UPPMAX, partially funded by the Swedish Research Council through grant agreement number 2018-05973.

## Author contributions

M.J. conceived the project. T.H., M.J., and M.M. designed experiments. T.H. performed experiments with assistance from I.L.V. and M.M.. I.L.V. prepared the labelled 4.5S RNA. T.H. performed data analysis. E.L. wrote code for visualization of data. All authors interpreted results. T.H. and M.J. wrote the paper with input from all other authors.

