## Supplementary Material for "Dynamic binding of the bacterial chaperone Trigger factor to translating ribosomes"

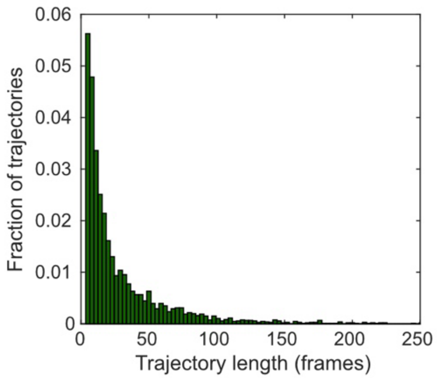


**Fig. S1. Trajectory length distribution for chromosomally expressed TF-Halo tracked with 5 ms camera exposure time**. Trajectories with at least 5 steps were included in the HMM fitting. n = 99,352 trajectory steps cumulated from 3 independent experiments.

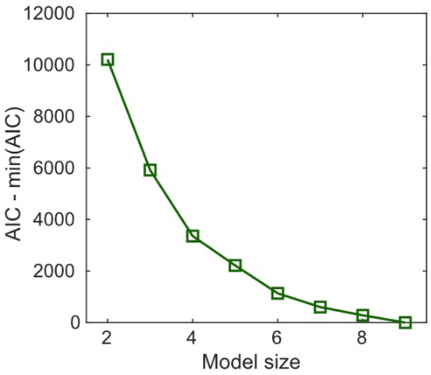


**Fig. S2. Akaike's information criterion (AIC) analysis of TF-Halo HMM-fitted models.** AIC scores, relative to the lowest one, for HMM fitting of trajectories of chromosomally expressed TF-Halo to different numbers of diffusion states (2-9). N = 99,352 trajectory steps cumulated from three independent experiments with 5 ms camera exposure time. The AIC score becomes smaller the more states are added.

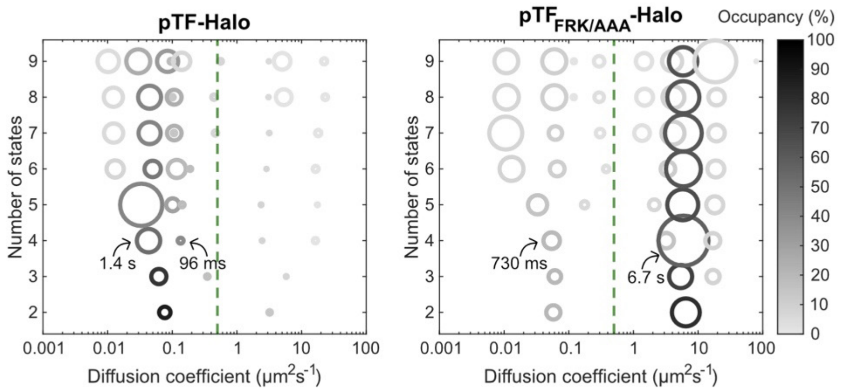


**Fig. S3. HMM models of TF-Halo and $TF_{FRK/AAA}$-Halo.** Leaky expression of TF-Halo and $TF_{FRK/AAA}$-Halo from plasmids in TF knockout cells. Circles are color-coded according to state occupancy and the area is proportional to the dwell time. Green lines mark a diffusion threshold of 0.5 $\mu m^2 s^{-1}$, separating free and RNC-bound TF-Halo. n = 83,546 and 75,585 trajectory steps from 3 and 5 independent experiments, respectively. Full model outputs are shown in Supplementary Data 6-7.

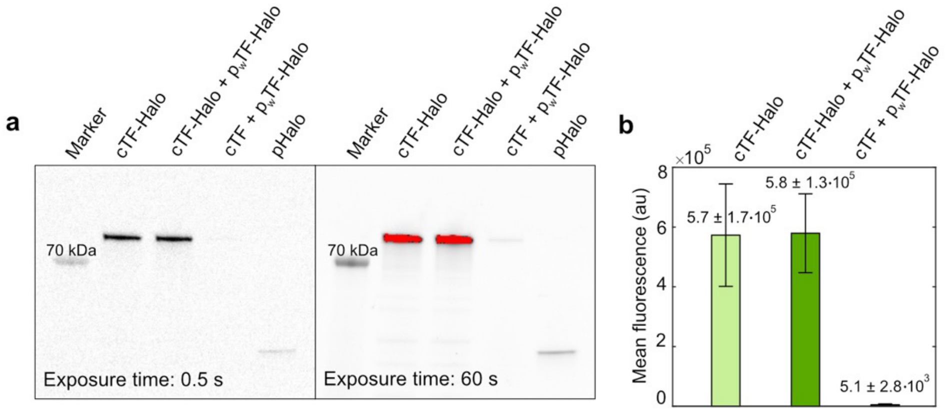


**Fig. S4. Levels of TF-Halo in strains used for SPT in Main Fig. 4. a** Example of SDS-PAGE gel imaged with green light to visualize fluorescence from TF-Halo labeled with JFX549. Levels of TF-Halo in each strain (cTF-Halo, cTF-Halo + $p_w$TF-Halo, cTF + $p_w$TF-Halo) were assessed by SDS-PAGE analysis of JFX549-labeled lysates. Strains were inoculated from glycerol stocks in Luria Broth (LB) and appropriate antibiotics and grown overnight at 37°C, 200 rpm. Overnight cultures were diluted 1:100 in LB and appropriate antibiotics and grown to $OD_{600} \approx 1.0$. 2 ml cell culture per 1 $OD_{600}$ was pelleted and resuspended in 100 µl Bacterial Protein Extraction Reagent (B-PER, Thermo Scientific), supplemented with cOmplete Mini EDTA-free protease inhibitors (Sigma Aldrich) and 0.3 µM JFX549, followed by incubation at room temperature for 10 min. Lysates were mixed 1:1 with 2x Laemmli Buffer (Bio-Rad) and B-mercaptoethanol. SDS-PAGE was run on 4-20% Mini-PROTEAN TGX pre-cast gels at 180V for 45 min. Gels were imaged with ChemiDoc MP (Bio-Rad) using green illumination for visualization of JFX549 signal. At 0.5 s exposure (left), there was strong signal for chromosomally expressed TF-Halo and only a weak signal from weak plasmid expression in wt background (cTF + $p_w$TF-Halo). For better visualization of the weak signal, an additional image was acquired with 60 ms exposure time (right), at which the stronger signals became saturated (red pixels). Hence, the 0.5 s exposure images were used to quantify the fluorescence signal intensity. **b** Mean fluorescence intensities from TF-Halo in each strain. Intensities were measured by densitometry from images with 0.5 s camera exposure time. Chromosomal expression of TF-Halo yields approximately 100-fold more TF-Halo than weak expression from the $p_w$TF-Halo plasmid. Mean intensities were calculated from n = 5 independent experiments and error bars show the standard deviation. Intensities from individual experiments are shown in Supplementary Data 23.

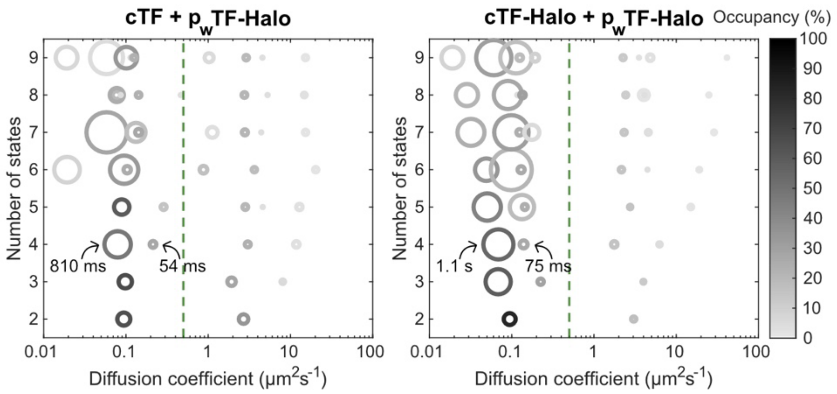


**Fig. S5. HMM models of TF-Halo in different background strains.** HMM fitting of 2-9 states of TF-Halo in the chromosomally tagged strain with additional weak plasmid expression (cTF-Halo + $p_w$TF-Halo) and in a wt strain with additional weak plasmid expression (cTF + $p_w$TF-Halo). n = 92,876 and 96,321 trajectory steps cumulated from 3 and 4 independent experiments for cTF-Halo + $p_w$TF-Halo and cTF + $p_w$TF-Halo, respectively. Circles are color-coded according to state occupancy and the area is proportional to the dwell time. Green lines mark a diffusion threshold of 0.5 μm$^2$s$^{-1}$, separating free and RNC-bound TF-Halo. Full model outputs are shown in Supplementary Data 12-13.

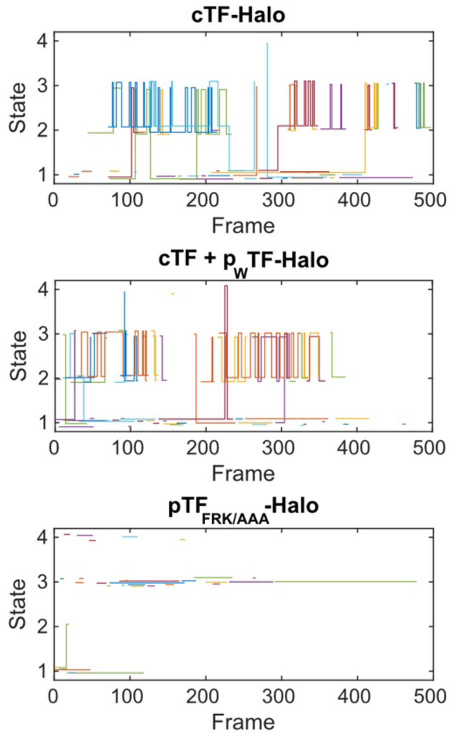


**Fig. S6. State transitions in a subset of TF-Halo and TF$_{FRK/AA}$-Halo trajectories fitted to 4-state models. a** Trajectories detected in cells from one SPT movie of chromosomally expressed TF-Halo (cTF-Halo), from low TF-Halo expression in the wt TF background (cTF + $p_w$TF-Halo) and from the FRK/AAA mutant. For TF-Halo, the most frequent transitions are between state 3 and 2, i.e., the free and RNC-sampling state. State 1 is target RNC binding. The FRK/AAA mutant, compromised in ribosome binding, does not exhibit the sampling behavior.

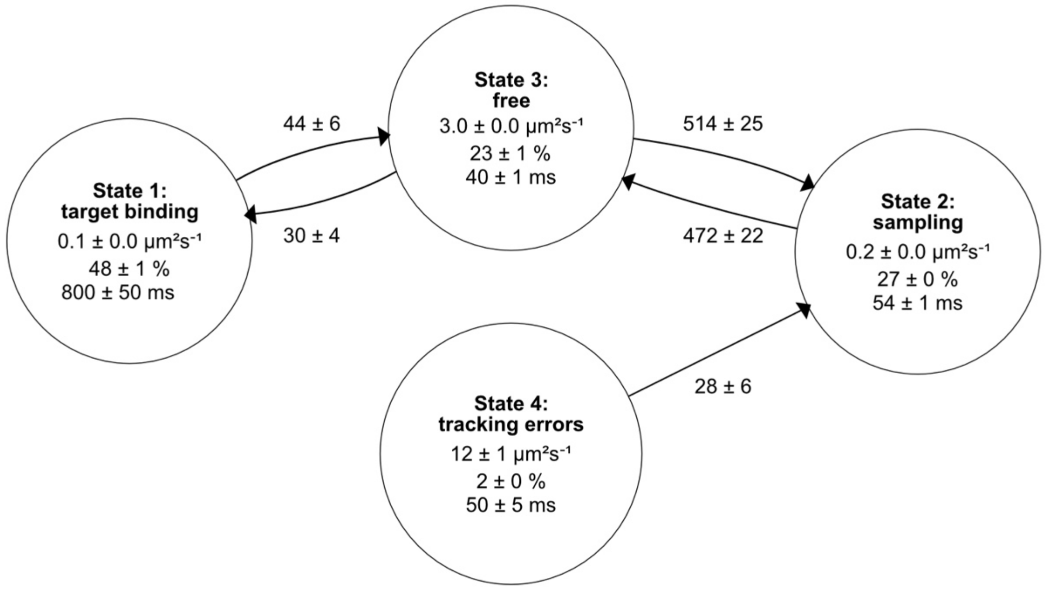


**Fig. S7. Fluxes of TF-Halo particles between states in HMM-fitted 4-state model.** Direction of fluxes are implied by the arrows. Fluxes are given as the percentage of the total population of TF particles transitioning between two states per second (%$s^{-1}$). Transitions between the free and RNC sampling state is predominant. Data comes from chromosomal expression of TF-Halo tracked at 5 ms, n = 99,352 trajectory steps cumulated from 3 independent experiments. Fluxes between states with 25 %$s^{-1}$ were excluded from the chart. Error bars for diffusion coefficients, occupancies and dwell times are bootstrap-estimated standard errors. For fluxes, the errors are propagated from the state occupancy and transition frequency bootstrap errors.

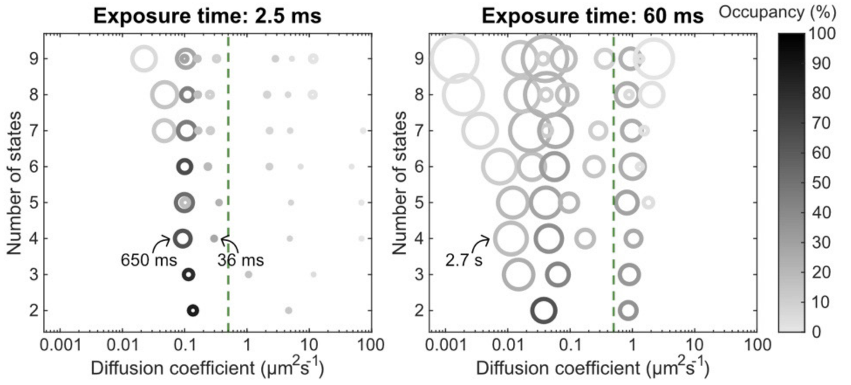


**Fig. S8. HMM models of chromosomally expressed TF-Halo using different camera exposure times.** With 2.5 ms exposure time, long and short RNC-bound states are resolved, similarly to the 5 ms data (main Fig. 2a). With 60 ms, the exposure time equals the average RNC sampling time, and consequently, the differences in dynamics between ribosome target-binding and sampling are lost. We note that for "free" TF-Halo with 60 ms exposures, the apparent diffusion becomes slower and dwell times become longer. This is attributed to averaging due to loss of transitions between the free and sampling states, as the average dwell time in the free state is < 60 ms, based on the 5 and 2.5 ms data. Circles are color-coded according to state occupancy and the area is proportional to the dwell time. Green lines mark a diffusion threshold of 0.5 $\mu m^2 s^{-1}$. n = 86,929 and 83,114 trajectory steps for 2.5 and 60 ms data, respectively, each cumulated from 3 independent experiments. For comparison, the x axes were set to [0.001, 100] for both the 2.5 and the 60 ms plot. However, model size 6 in 2.5 ms tracking contains a low occupancy (< 0.1%) slow artefact state with D = $10^{-6}$ $\mu m^2 s^{-1}$, not included in the plot. Full model outputs are shown in Supplementary Data 14-15.

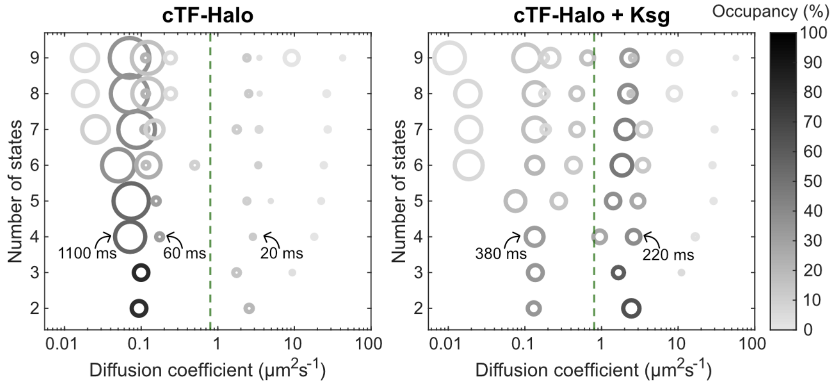


**Fig. S9. HMM models of chromosomally expressed TF-Halo in cells treated with 2 mg/ml Kasugamycin (Ksg).** The cTF-Halo plot (left) is the same data as shown Main Fig. 2a and is displayed here as a reference for the effect of Ksg treatment (right). Upon Ksg treatment, dwell times in the free states become longer for all model sizes. The distinction between target RNC binding and sampling becomes evident only in models with 7-9 states and the sampling state has reduced occupancy (roughly 6%, Supplementary Data 16). Together, this suggests that TF-Halo performs less sampling in Ksg-treated cells. Thus, TF-Halo does not sample free 50S subunits, but rather requires interactions both with the ribosomal surface and a nascent chain. Circles are color-coded according to state occupancy and the area is proportional to the dwell time. Green lines mark a diffusion threshold of 0.8 $\mu m^2 s^{-1}$. n = 99,352 and 85,146 trajectory steps for untreated and Ksg-treated cells, respectively, each cumulated from 3 independent experiments. Full model outputs are shown in Supplementary Data 4 and 16.

*Table S1: List of primer sequences*

| Name | Sequence, 5' -> 3' |
|---|---|
| tig_HaloIns_F | aaagaaaccactttcaacgagctgatgaaccagcaggcgggcgcagaaatcggtactggc |
| tig_del_R | acgggcctttgtgcgaatttagcgcgttatgctgcgtaaagtgtaggctggagctgcttc |
| pQE_SDw_F | tttcacacagaattcattaaagtttcttaattaactatgcaagtttcagttgaaac |
| p124_R | atggctgtaagtattcgccgcaagggataaatgtcgatttctcgaggtgaagacgaaagg |

### Supplementary Movie 1

Microscopy data of TF-Halo diffusion in live *E coli*. TF-Halo was expressed from the chromosome and labeled with JFX549. The movie (middle panel) was acquired with 5 ms camera exposure time and 3 ms illumination (546 nm) per image. For analysis, movies were aligned with cell outlines (segmented based on phase contrast images, bottom panel), and trajectories of single TF-Halo particles were built using the uTrack algorithm and HMM-fitted to a 2-state diffusion model with state 1 corresponding to slow diffusion and state 2 to fast diffusion (top panel). To reduce the risk of errors in the trajectory building, trajectories were recorded when there was only one fluorescent dot detected in a cell. Playback speed is 20 frames per second, i.e, 10 times slower than reality.

### Supplementary Movie 2

Microscopy data of $TF_{FRK/AAA}$-Halo diffusion in live *E coli*. $TF_{FRK/AAA}$-Halo was expressed from an IPTG-inducible plasmid at leaky expression level in a TF knockout strain and labeled with JFX549. The movie (middle panel) was acquired with 5 ms camera exposure time and 3 ms illumination (546 nm) per image. For analysis, movies were aligned with cell outlines (segmented based on phase contrast images, bottom panel), and trajectories of single TF-Halo particles were built using the uTrack algorithm and HMM-fitted to a 2-state diffusion model with state 1 corresponding to slow diffusion and state 2 to fast diffusion (top panel). To reduce the risk of errors in the trajectory building, trajectories were recorded when there was only one fluorescent dot detected in a cell. Playback speed is 20 frames per second, i.e, 10 times slower than reality.

### Supplementary Movie 3

Example of a TF-Halo trajectory HMM-fitted to a 4-state model displaying the RNC sampling behavior, i.e., frequent transitions between freely diffusing (state 3) and a slow-diffusing state (state 2). TF-Halo was expressed from the chromosome and labeled with JFX549. The movie was acquired with 5 ms camera exposure time and 3 ms illumination (546 nm) per image. State 1 corresponds to long target RNC binding, state 2 to shorter RNC sampling, and state 3 to free TF-Halo. Playback speed is 20 frames per second, i.e, 10 times slower than reality.

### Supplementary Movie 4

Example of a TF-Halo trajectory HMM-fitted to a 4-state model displaying long RNC binding. TF-Halo was expressed from the chromosome and labeled with JFX549. The movie was acquired with 5 ms camera exposure time and 3 ms illumination (546 nm) per image. State 1 corresponds to long target RNC binding, state 2 to shorter RNC sampling, and state 3 to free TF-Halo. Playback speed is 20 frames per second, i.e, 10 times slower than reality.

### Supplementary Movie 5

Example of a $TF_{FRK/AAA}$-Halo trajectory HMM-fitted to a 4-state model. $TF_{FRK/AAA}$-Halo was expressed from an IPTG-inducible plasmid at leaky expression level in a TF knockout strain and labeled with JFX549. The movie was acquired with 5 ms camera exposure time and 3 ms illumination (546 nm) per image. State 1 corresponds to long target RNC binding, state 2 to shorter RNC sampling, and state 3 to free TF-Halo. Playback speed is 20 frames per second, i.e, 10 times slower than reality.
